# Numerical analysis of flow anisotropy in rotated-square deterministic lateral displacement devices at moderate Reynolds number

**DOI:** 10.1101/2023.10.02.560085

**Authors:** Calum Mallorie, Rohan Vernekar, Benjamin Owen, David W. Inglis, Timm Krüger

## Abstract

Deterministic lateral displacement (DLD) is a microfluidic method for accurately separating particles by size or deformability. Recent efforts to operate DLD devices in the inertial, rather than in the Stokes, flow regime have been hindered by a loss of separation efficiency and difficulty predicting the separation behaviour. One factor contributing to these problems is the onset of inertia-induced flow anisotropy where the average flow direction does not align with the direction of the pressure gradient in the device. We use the lattice-Boltzmann method to simulate two-dimensional flow through a rotated-square DLD geometry with circular pillars at Reynolds number up to 100 for different gap sizes and rotation angles. We find that anisotropy in this geometry is a non-monotonous function of Reynolds number and can be positive or negative. This finding is in contradiction to the naive expectation that inertia would always drive flow along principal direction of the pillar array. Anisotropy tends to increase in magnitude with gap size and rotation angle. By analysing the traction distribution along the pillar surface, we explain how the change of the flow field upon increasing inertia leads to the observed trends of anisotropy. Our work contributes to a better understanding of the inertial flow behaviour in ordered cylindrical porous media, and might contribute to improved DLD designs for operation in the inertial regime.

## I. INTRODUCTION

Particle separation based on physical properties is an essential process in medical diagnostics and drug development. Microfluidic techniques, including deterministic lateral displacement (DLD), have gained prominence due to their high size resolution and throughput compared to traditional methods like filtration and centrifugation [1]. DLD was originally proposed in 2004 by Huang *et al*. [2] and has been successfully applied to the separation of various biological particles by their mechanical phenotype [3, 4], with a size resolution down to 10 nm [2].

DLD operates by dividing a laminar fluid flow into flow lanes that are bounded by stagnation streamlines which originate and terminate at the surface of pillars arranged in a periodic array of obstacles [5], see Fig. 1. To leading order, particles follow streamlines and, therefore, remain in the same flow lane, unless forced out by contact with an obstacle. If a particle is larger than a critical radius *r*_c_, determined by the width of the flow lane adjacent to a pillar, contact with a pillar surface forces the hydrodynamic centre of the particle into a neighbouring flow lane. This process occurs at each subsequent pillar, leading to lateral migration across streamlines. Particles smaller than *r*_c_ remain in their flow lane and travel in the same direction as the bulk fluid flow. The different modes of transport through the DLD device have different angles of displacement relative to the flow field and different particles can be collected at separate downstream outlets.

**FIG. 1.**
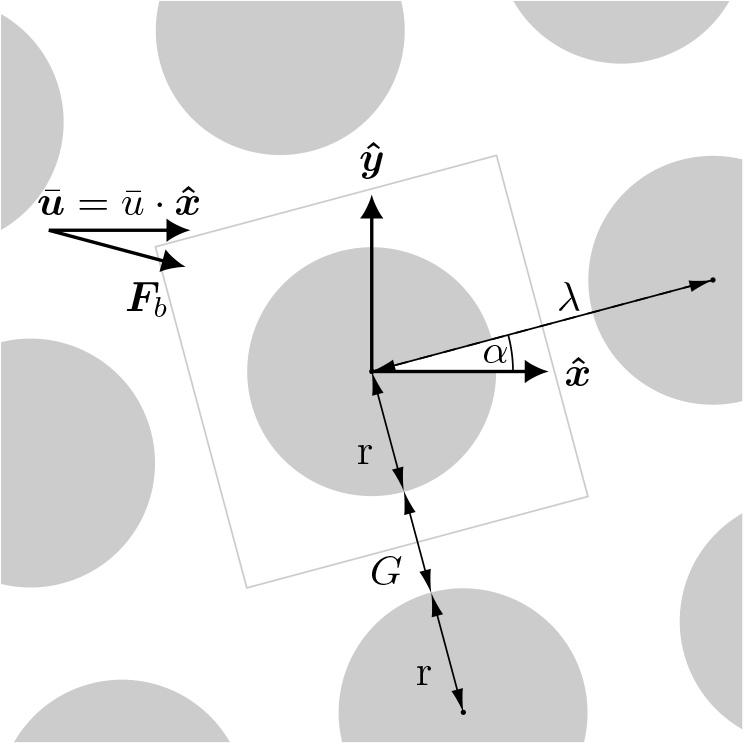
Illustration of the rotated-square array. The circular pillars of radius *r* lie on the nodes of a square lattice with centre-to-centre distance *λ*. The gap size between pillars is *G* = *λ−* 2*r*. The array is rotated at an angle *α* relative to the coordinate axes. The average velocity of the fluid 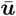 is imposed to be parallel to the basis vector 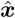 which indicates the axis of the DLD device. The body force density driving the flow is denoted ***F***_b_. Simulations are carried out over one unit cell of the array with periodic boundary conditions, highlighted by the grey box.

DLD has been shown to be highly effective in the Stokes regime, with separation efficiencies approaching 100% in some cases [6, 7]. The behaviour of the devices in this regime has been well characterised, with predictive empirical relations [8] and design heuristics [9] developed. However, operating devices in the Stokes regime limits the sample throughput due to the necessarily low flow rates.

Operating DLD devices with higher flow rates requires larger flow speed, which leads to the onset of inertial effects. DLD devices operating in the inertial regime show a reduction of the critical size [10–12], which is useful to prevent clogging. However, inertia also leads to additional effects, such as breakdown of the separation efficiency [13], loss of symmetry in the flow field [12, 14], and increased influence of the particle-fluid density difference [15].

In some examples of DLD operating at high flow rates, there is evidence that the average flow direction tilts to align with the array inclination as the flow rate is increased [16, 17]. A likely cause of this problem is anisotropic fluid conduction in the array [16]. Anisotropy is the tendency of the fluid to deviate from the direction of the applied pressure gradient. Anisotropy can cause changes to the average flow direction, which changes the critical size in the array in complex ways and causes a breakdown of separation performance [18, 19]. DLD devices with circular pillars arranged in a rotated-square layout (Fig. 1) have zero anisotropy in the Stokes regime [20], and are therefore recommended as best practice [21]. Under the influence of inertia, however, rotated-square layouts have been shown to exhibit anisotropy [22]. The mechanisms leading to the increase in anisotropy with fluid inertia in DLD geometries should be better understood to enable improved designs of inertial DLD devices.

Flows though DLD-like cylindrical pillar arrays have been historically investigated in the field of ordered porous media studies [23, 24], which have numerous industrial [25, 26], biological [27], and geological [28] applications. Most of these studies choose pillar array alignment of either 0° or 45° to the driving pressure gradient, ensuring flow symmetry in steady state, with a strong interest in measuring the flow resistance (or permeability) of the arrays [29–34]. As far as we are aware, only Koch and Ladd [22] have examined the effect of fluid inertia on flow permeability of inclined cylindrical pillar arrays. They found that rotated-square arrays show anisotropic permeability due to fluid inertia. Remarkably, they observed that the anisotropy can become negative for certain configurations, meaning that the flow chooses to tilt against the inclination of the pillar array, due to fluid inertia. Koch and Ladd [22] did not explain the physical reasons for this counter-intuitive behaviour.

In this work, we use a two-dimensional (2D) lattice-Boltzmann method to simulate inertial flow in rotated-square DLD devices. We investigate the emergence of anisotropy as a function of Reynolds number, gap size and array inclination angle. Our results confirm that anisotropy is a non-monotonous function of Reynolds number and can be positive or negative. The magnitude of anisotropy generally increases with gap size and array inclination. By analysing the distribution of the traction vector along the pillar surface we explain the observed behaviour of the anisotropy.

## II. MODEL AND METHODS

### A. Physical and mathematical model

We investigate the flow of a Newtonian, incompressible fluid through a rotated-square DLD device in 2D (Fig. 1). We consider a portion of the device far from the walls that confine the DLD device. Circular pillars of radius *r* are arranged in a periodically repeating square array with centre-to-centre distance *λ*. The minimum gap size between pillar surfaces is *G* = *λ−* 2*r*. The array is tilted at an angle *α* counterclockwise with respect to the device axis. The no-slip condition holds at the surface of the pillars, and periodic boundary conditions are applied to the simulation domain.

We define the unit basis vectors 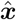 and 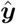 to point tangentially and orthogonally to the axis of the DLD device. The average flow direction 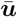 is aligned with the *x*-axis,

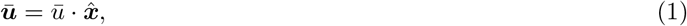

as would be expected in a real-world device. The flow is driven by a body force density (force per unit area in 2D) ***F***_b_ whose direction is set such that Eq. (1) holds [20]. We only consider cases for which the flow field becomes steady after some time, which, for given values of *G* and *α*, happens at sufficiently low Reynolds number.

### B. Drag and lift forces and flow anisotropy

The fluid stress tensor ***σ***, including inertial, pressure and viscous contributions, can be expressed using index notation as

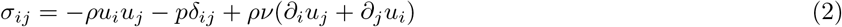

where *ρ* is density, *ν* is kinematic viscosity, *u*_*i*_ is the *i*-component of the fluid velocity, *p* is pressure, and *δ*_*j*_ is the Kronecker symbol. On the pillar surface *S*, the inertial contribution, *ρu*_*i*_*u*_*j*_, vanishes due to the no-slip condition:

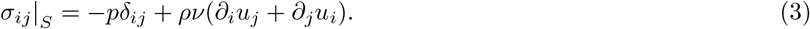

The traction vector ***t*** at any point on *S* can be calculated by projecting the stress tensor onto the surface normal vector 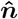 which, by definition, points into the fluid:

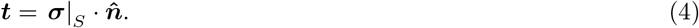

Note that ***t*** is defined in such a way that the traction vector is the force exerted by the fluid on the pillar surface, rather than the other way around.

At steady state, the momentum balance dictates that the integral of the traction vector over the surface *S* of one pillar is equal to the integral of the body force driving the fluid flow over the area *A* of one unit cell of the array:

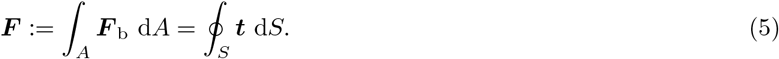

The local drag and lift force densities acting on a pillar surface element d*S* are the components of the traction vector pointing in the direction of, and orthogonally to, the average flow direction, respectively:

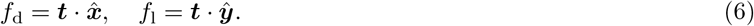

The total force acting on the pillar can be decomposed into net drag and lift forces, 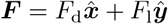, where

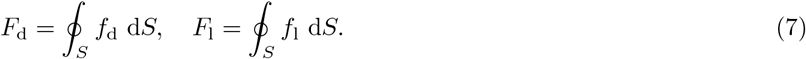

Note that *F*_d_ is always positive and *F*_l_ can be positive, negative or zero.

A flow configuration for which *F*_l_ = 0 holds for any angle *α* is called isotropic: the average flow 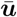 and the total force vector ***F*** _b_ are always parallel, and no lateral flow deflection occurs. The prime example for an isotropic flow is the Stokes flow (zero Reynolds number) over a square array of circular obstacles [20]. Isotropy can be broken by choosing different pillar shapes, pillar arrangements or by having a finite Reynolds number [20, 22, 35]. We define flow anisotropy as the dimensionless ratio of lift over drag:

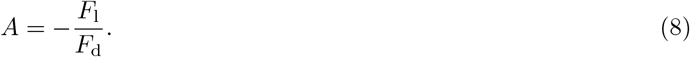

The negative sign is to match the convention established in previous work [20]. Positive anisotropy, *A >* 0, indicates that the flow would be deflected in a counterclockwise fashion (along the positive *y*-axis). However, in a real device, fluid would not be able to accumulate at the side walls due to incompressibility of the fluid, and instead a lateral pressure gradient would build up that maintains the average flow along the *x*-axis. In our model, this pressure gradient is captured by the *y*-component of the body force density ***F*** _b_ which is obtained in such a way that Eq. (1) holds at any time. The resulting force components *F*_d_ and *F*_l_ then determine *A*.

For further analysis, we also define the contributions to flow anisotropy coming from the normal and shear components of the traction vector, respectively:

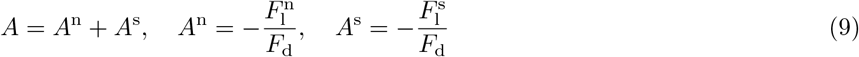

where

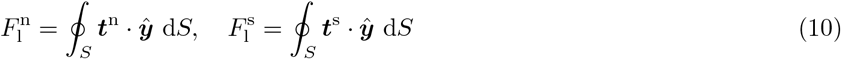

where

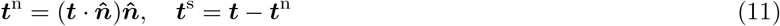

are the normal and shear components of the traction vector. While *A*^s^ is determined by the viscous shear stress at the pillar surface, *A*^n^ follows from the viscous normal stress and the pressure at the pillar surface.

### C. Dimensionless numbers

Apart from the non-dimensionalised gap size *G*^⋆^ = *G/λ* and the array tilt angle *α*, the problem is fully characterised by the Reynolds number *Re*. We define *Re* by the average flow velocity 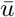 in the fluid domain excluding the pillars, the pillar diameter 2*r*, and the kinematic viscosity *ν*, similarly to Koch and Ladd [22]:

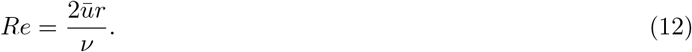

It is also useful to define the packing fraction of the pillars as 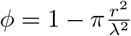 . On a macroscopic level, the converged flow is fully described by the resulting anisotropy *A* (and its contributions *A*^n^ and *A*^s^).

### D. Numerical model and simulation parameters

The 2D fluid flow is solved using the lattice-Boltzmann (LB) method on a D2Q9 lattice [36, 37]. We use the BGK collision operator [38] and the Guo forcing scheme [39]. The simple bounce-back method [40] recovers the no-slip boundary condition at the surface of the pillar; this leads to a staircase approximation for all curved boundary surfaces. The kinematic viscosity is set by the lattice-Boltzmann relaxation time, *τ* [36]:

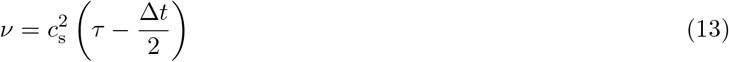

where 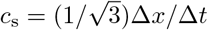 is the speed of sound, Δ*x* is the lattice spacing and Δ*t* is the time step.

The simulation domain is one periodic unit cell of the DLD array, shown by the grey box in Fig. 1. The simulation frame is converted to the physical frame by a clockwise rotation of *α*.

To impose the average direction of the velocity field 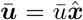 and its desired magnitude 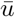, the body force density ***F*** _b_ is determined iteratively. Only steady cases are considered here, and simulations are stopped when the flow has converged with a tolerance of 10^*−*8^ calculated from the *L*_2_ norm of the difference of the flow fields at times *t* and *t−* 1000Δ*t*.

To ensure the fluid flow and obstacle surface shape are approximated with sufficient accuracy, we carried out convergence studies (Fig. 2a). We found that a domain size of 200 *×* 200 grid points is sufficient. In the following, all reported results have been obtained on a lattice with 200 *×* 200 grid points, corresponding to *λ* = 200Δ*x*.

**FIG. 2.**
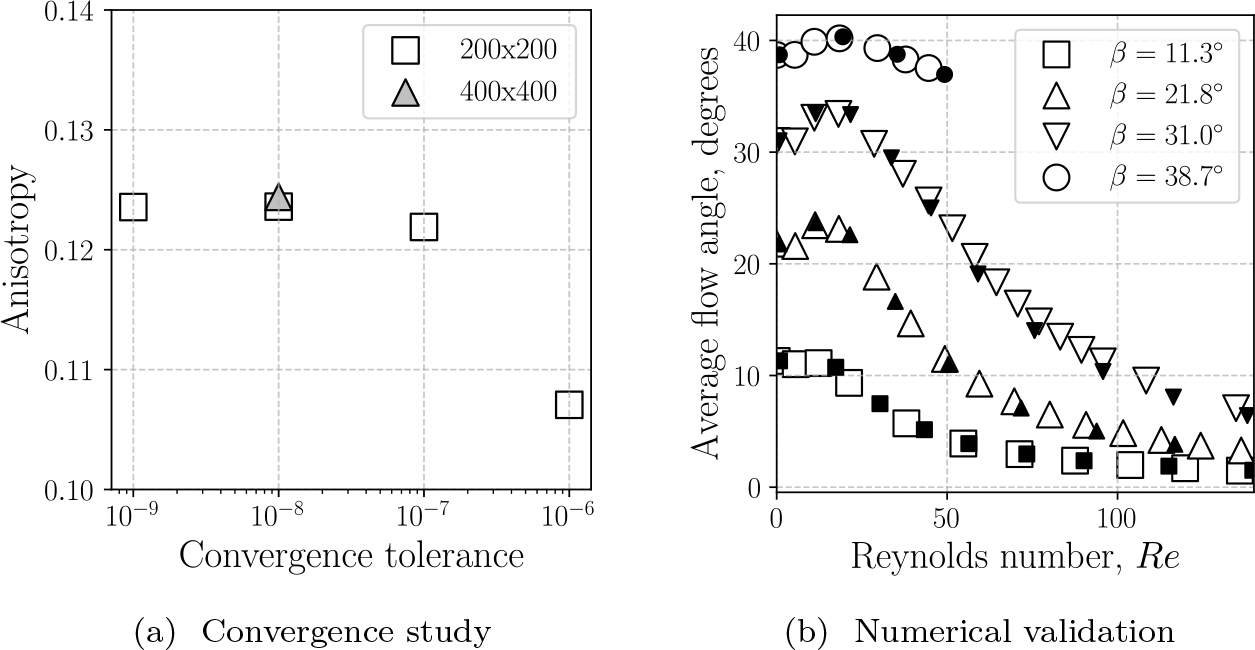
(a) Study of convergence tolerance and grid independence with *G*^⋆^ = 0.5, *Re* = 50. Doubling the resolution to 400 × 400 grid points increases the anisotropy by 0.65% at a convergence tolerance of 10^−8^ between times *t* and *t −*1000Δ*t*. (b) Average flow direction relative to the array orientation at *G*^⋆^ = 0.51 against *Re*, for simulations with the flow allowed to freely develop (body force at an angle *β* with respect to 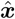). In black are the results obtained by Koch and Ladd [22].

### E. Obtaining the fluid stress on the pillar surface

Calculation of the traction vector acting on the pillar surface by Eq. (4) requires knowledge of the fluid stress tensor on the surface. We use an interpolation-extrapolation scheme to obtain the fluid stress tensor on the pillar surface.

We discretise the cylinder surface as a series of *N* evenly spaced points with separation ≈ Δ*x* such that 2*πr ≈ N* Δ*x*. For each of the *N* surface poins, we define two points along the surface normal vector at a distance of Δ*r* and 2Δ*r* where we use Δ*r* = 2Δ*x*. Bilinear interpolation is used to find the fluid stress tensor at these two points using the four fluid nodes delineating the fluid voxel in which the interpolation point resides. Linear extrapolation from the two points is used to determine the fluid stress tensor at each of the *N* surface points. The fluid stress tensor is then smoothed by computing a moving average along the cylinder surface to reduce noise from the interpolation, with averages taken over five points for the normal components of the stress and two points for the shear component.

## III. RESULTS AND DISCUSSION

### A. Validation: comparison with earlier work

To validate our simulations, we compare with the results obtained by Koch and Ladd [22] who investigated the behaviour of fluid flow over periodic square arrays of cylinders with *G*^⋆^ = 0.51, at moderate inertia. Koch and Ladd did not impose a specific flow direction (as we do in the present study), but rather imposed a body force with fixed direction *β* on an unrotated lattice (*α* = 0) and then measured the resulting average flow direction. Using the same set-up, we obtain similar results, as shown in Fig. 2b. Note that for the forcing angle *β* = 38.7°, Koch and Ladd found unsteady flows at *Re >* 50, which we also confirm. However, we focus on steady flows in the remainder of the present study.

The parameters for all remaining simulations are summarised in Tab. I.

**TABLE I.** Simulation parameters and associated dimensionless numbers. All quantities are expressed in simulation units (Δ*x* = 1 and Δ*t* = 1).

| $Re$ [-] | $G^*$ [-] | $\alpha$ [°] | $\tau$ | $\nu$ | $\bar{u}$ |
| --- | --- | --- | --- | --- | --- |
| [0.1, 100] | 0.2, 0.4, 0.6 | 1, 3, 5 | 0.55 | 1/60 | [$10^{-5}$, 0.02] |

### B. Flow field and velocity profiles

Fig. 3 shows streamlines and velocity magnitudes for simulations with *G*^⋆^ = 0.2, 0.4 and 0.6, *α* = 5°, and *Re* = 0.1 and 100. While the flow field is point symmetric about the centre of the pillar for small *Re*, a recirculating wake is visible downstream from the pillar at higher *Re*, becoming more pronounced with increasing *G*^⋆^. The fluid speed is much lower in the recirculation zones, with the majority of fluid flow concentrated in the horizontal gap between pillars, and in a high velocity stream flowing downwards (negative *y*-direction) between the recirculation zone and the adjacent downstream pillar.

**FIG. 3.**
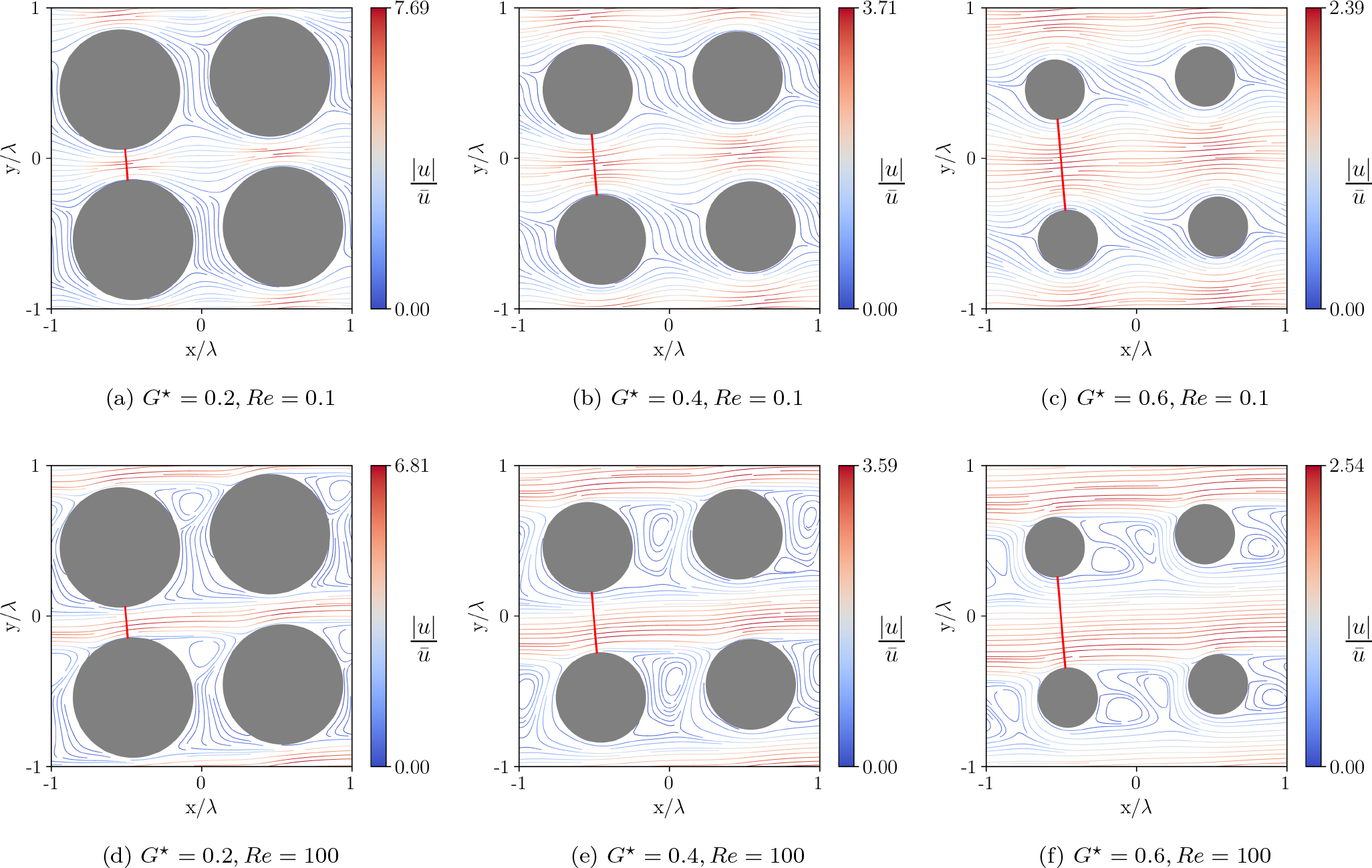
Streamlines coloured by the normalised velocity magnitude (see right colour bar) for different *G*^⋆^ at *α* = 5*°*. At low *Re* (a–c), the flow field is symmetrical and attached to pillar surfaces. At higher *Re* (d–f), a recirculating wake is visible downstream from the pillar. The red line illustrates the path over which the velocity profiles in Fig. 4 are sampled. Note that four periodic unit cells (and therefore four pillars) are shown for visual clarity.

Fig. 4 depicts the velocity profiles for *α* = 5° and different values of *Re* and *G*^⋆^ in the gap between pillars as indicated by the red lines in Fig. 3. While the velocity profile is symmetric and nearly parabolic for *Re* = 0.1 (near Stokes regime) for all studied values of *G*^⋆^, increasing *Re* leads to the velocity profile being skewed and pushed against the bottom of the confining gap (in negative *y*-direction), thus, the flow speed above the pillar is higher than below. This skewing effect becomes stronger with increasing *G*^⋆^ and *α* (data for other angles not shown) and is caused by the inertia of the fluid which is forced to flow, on average, along the *x*-axis.

**FIG. 4.**
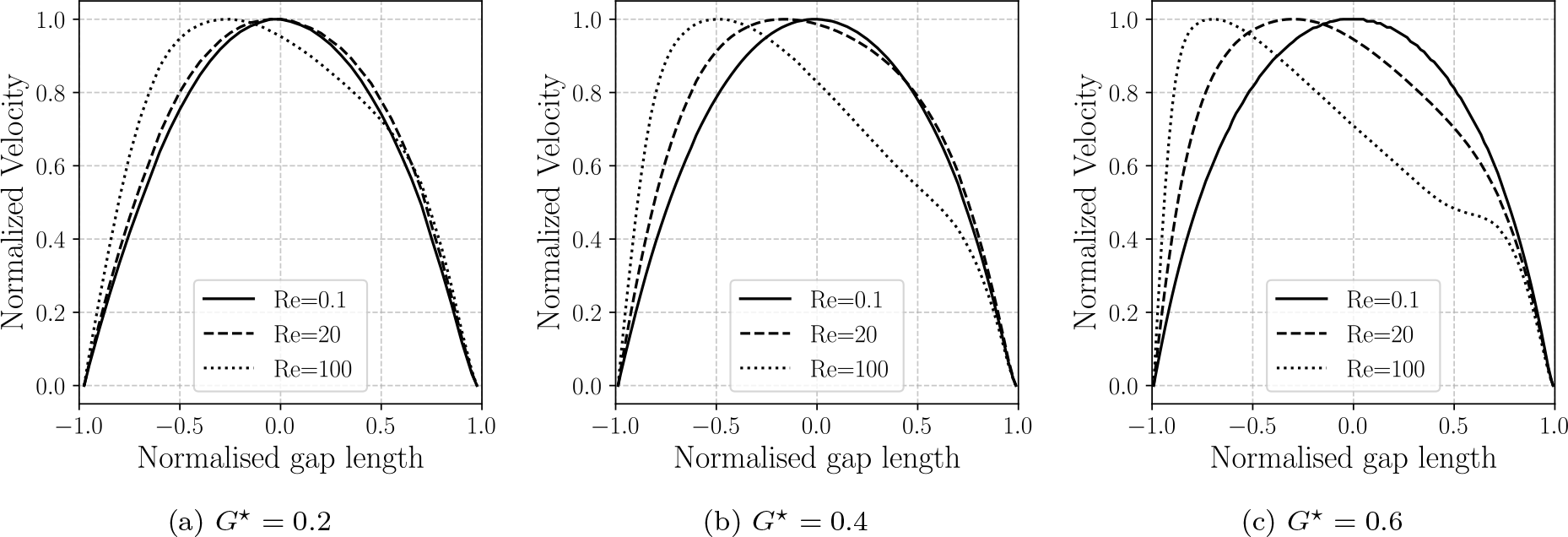
Velocity profiles along the vertical centre-to-centre line (shown as red line in Fig. 3) for *α* = 5*°* and different values of *G*^⋆^ and *Re*. The *x*-axis is normalised such that the top side of the pillar corresponds to −1 and the bottom side of the neighbouring pillar to +1. The *y*-axis is normalised by the value of the maximum velocity magnitude.

### C. Flow anisotropy

Fig. 5 illustrates the anisotropy *A* as a function of *Re* for various values of reduced gap size *G*^⋆^ and array inclination angle *α*. As expected, anisotropy tends towards zero for *Re →* 0 across all configurations. For *Re >* 5, anisotropy becomes significant as inertial effects start to play an important role.

**FIG. 5.**
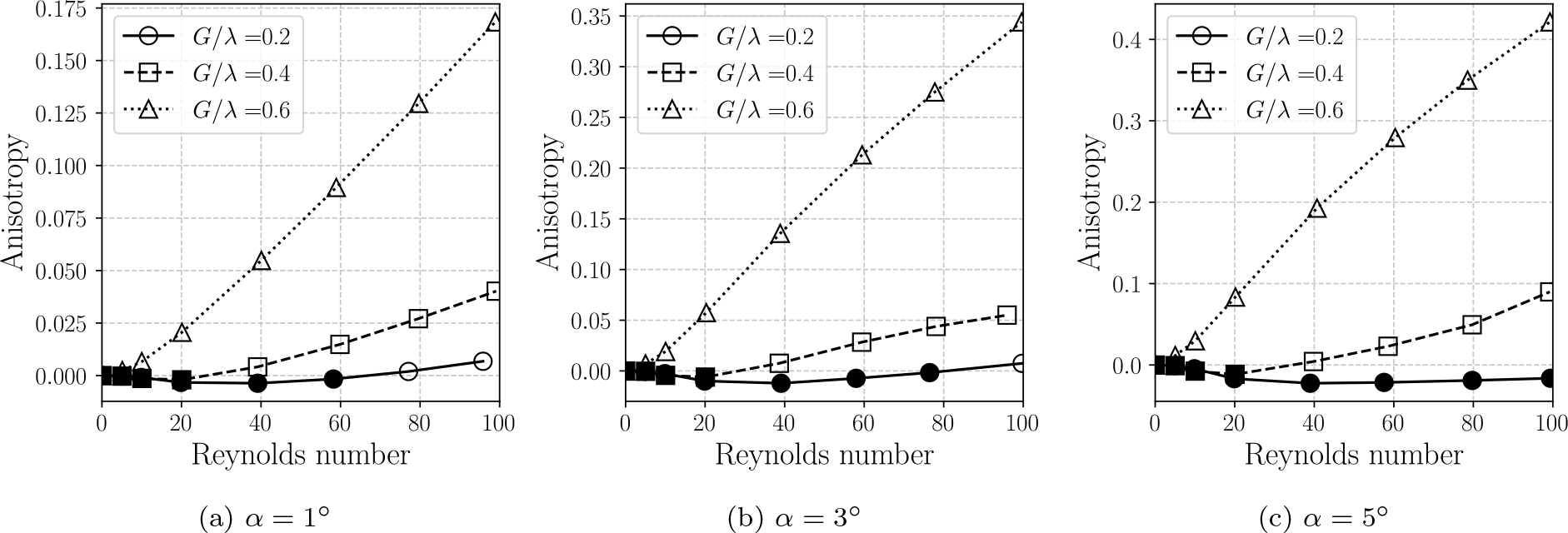
Anisotropy *A* for different values of *G*^⋆^, *α*, and *Re*. Results are grouped by the value of *α* to highlight the similarity of the results for different values of *α*. Note that the scale of the *y*-axis changes between panels. Points for which the anisotropy is negative are marked with filled shapes.

The behaviour of anisotropy also depends on gap size. For the largest gap size, *G*^⋆^ = 0.6, anisotropy grows strongly with *Re*, indicating that the fluid prefers to flow along the principle axis of the array. The magnitude of anisotropy also tends to grow with increasing gap size. This is consistent with expectations, as the drag force at a given Reynolds number will be smaller for larger gaps. If the lift were constant, the magnitude of the anisotropy would increase with decreasing gap size according to Eq. 8.

For smaller gaps (*G*^⋆^ = 0.4 and *G*^⋆^ = 0.2), anisotropy exhibits complex behaviour. As noted by Koch and Ladd in 1997 [22], anisotropy first becomes negative before increasing again at larger *Re*. Particularly for the smallest gap size, *G*^⋆^ = 0.2, and the largest array inclination angle, *α* = 5°, the anisotropy is negative for all Reynolds numbers studied.

Our results further reveal that the reversal of the direction of the lift force at intermediate *Re* occurs even at much smaller angles than those investigated by Koch and Ladd.

The contributions to the anisotropy from the normal and shear components of the traction vector are shown in Fig. 6(a–c) and Fig. 6(d–f), respectively. It is evident from these data that both contributions are of similar magnitude, hence neither can be neglected when analysing the mechanisms that lead to the overall behaviour. The data further demonstrate that shear forces always contribute positively to the total anisotropy and increase monotonically with gap size. Thus, as seen in Fig. 6(a–c), normal contributions are the sole source of the negative anisotropy observed for smaller gap sizes. There appear to be at least two competing *Re*-dependent mechanisms that determine the contribution of normal forces to the anisotropy, with the negative contributions dominating at intermediate values of *Re*, thus leading to the minimum, and negative values, of *A*. To understand these observations better, we need to look into the distribution of traction and lift along the pillar surface.

**FIG. 6.**
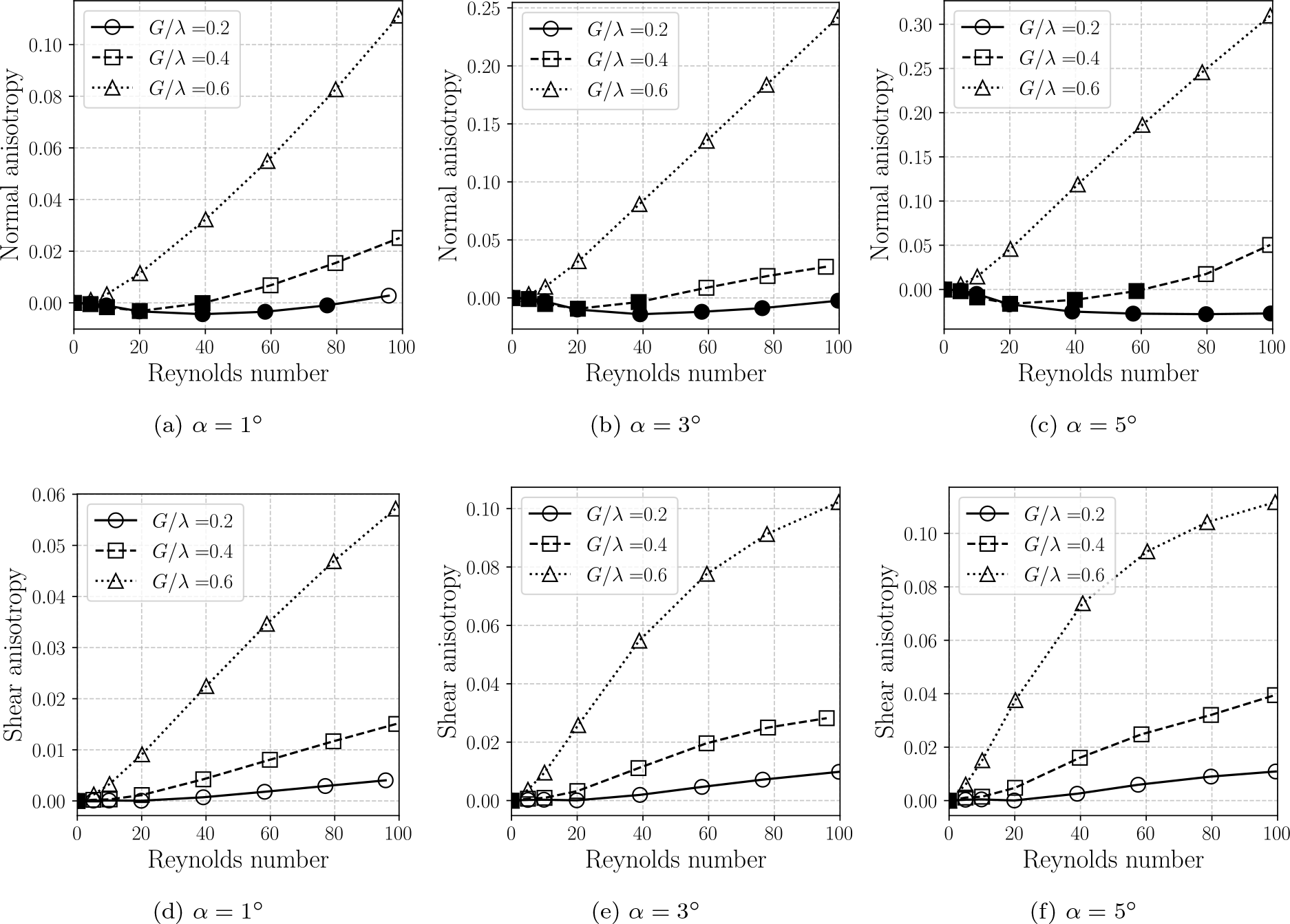
(a–c) Normal and (d–f) shear contributions to anisotropy for the simulations shown in Fig. 5. Points for which the anisotropy is below zero are marked with filled shapes.

### D. Link between local lift profiles and anisotropy

In order to understand the mechanisms leading to the onset of anisotropy and its dependency on *Re* for a given geometry (*G*^⋆^ and *α*), we investigate the local forces acting on the pillar surface. Apart from the upstream (*x <* 0), downstream (*x >* 0), bottom (*y <* 0) and top (*y >* 0) sides of the pillar, we define four quadrants for convenience (see annotation of Fig. 7a):

- Bottom left (BL): *x <* 0, *y <* 0
- Top left (TL): *x <* 0, *y >* 0
- Top right (TR): *x >* 0, *y >* 0
- Bottom right (BR): *x >* 0, *y <* 0

**FIG. 7.**
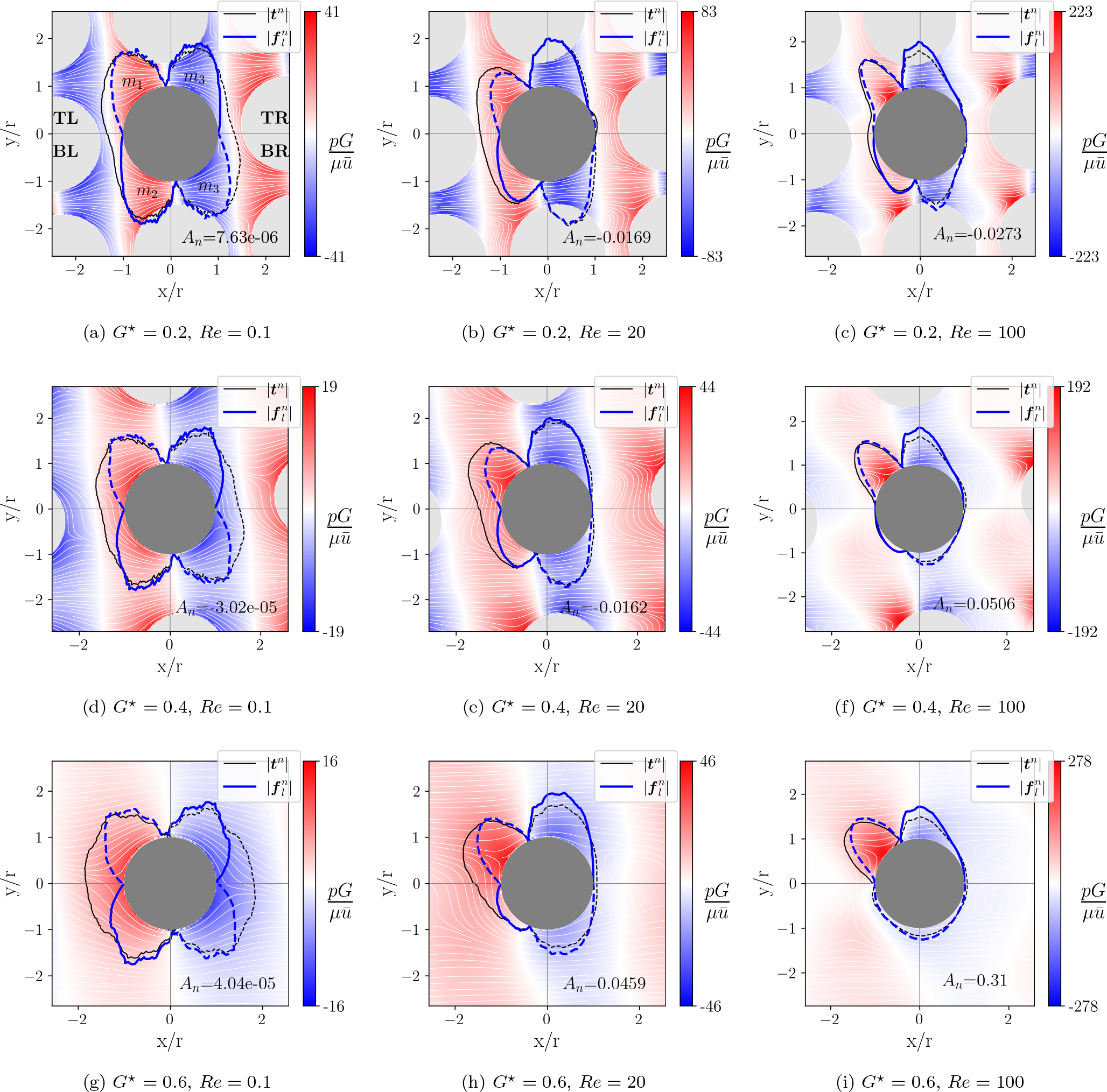
Polar representation of the normal contribution to the traction (blue) and lift (black) along the pillar surface for (a–c) *G*^⋆^ = 0.2, (d–f) *G*^⋆^ = 0.4 and (g–i) *G*^⋆^ = 0.6 at *α* = 5*°* for different values of *Re*. The average flow direction is along the *x*-axis. The traction and lift distributions are normalised by their maximum values. Positive values are shown as a solid line, negative values as a dashed line. The value of the normal contribution to the anisotropy is reported in each panel. Panel (a) also contains labels of the quadrants, and the labels for mechanisms associated with each region of the lift distribution, as referred to in section III D. The normalised pressure field is shown as a blue-white-red colour gradient.

Fig. 7 and Fig. 8 show the normal and shear contributions, respectively, to the local traction and lift force along the pillar surface for different values of *G*^⋆^ and *Re* at *α* = 5°. Note that the normal contribution to the local traction and lift is dominated by the pressure forces; the normal component of the viscous stress is negligible (data not shown).

**FIG. 8.**
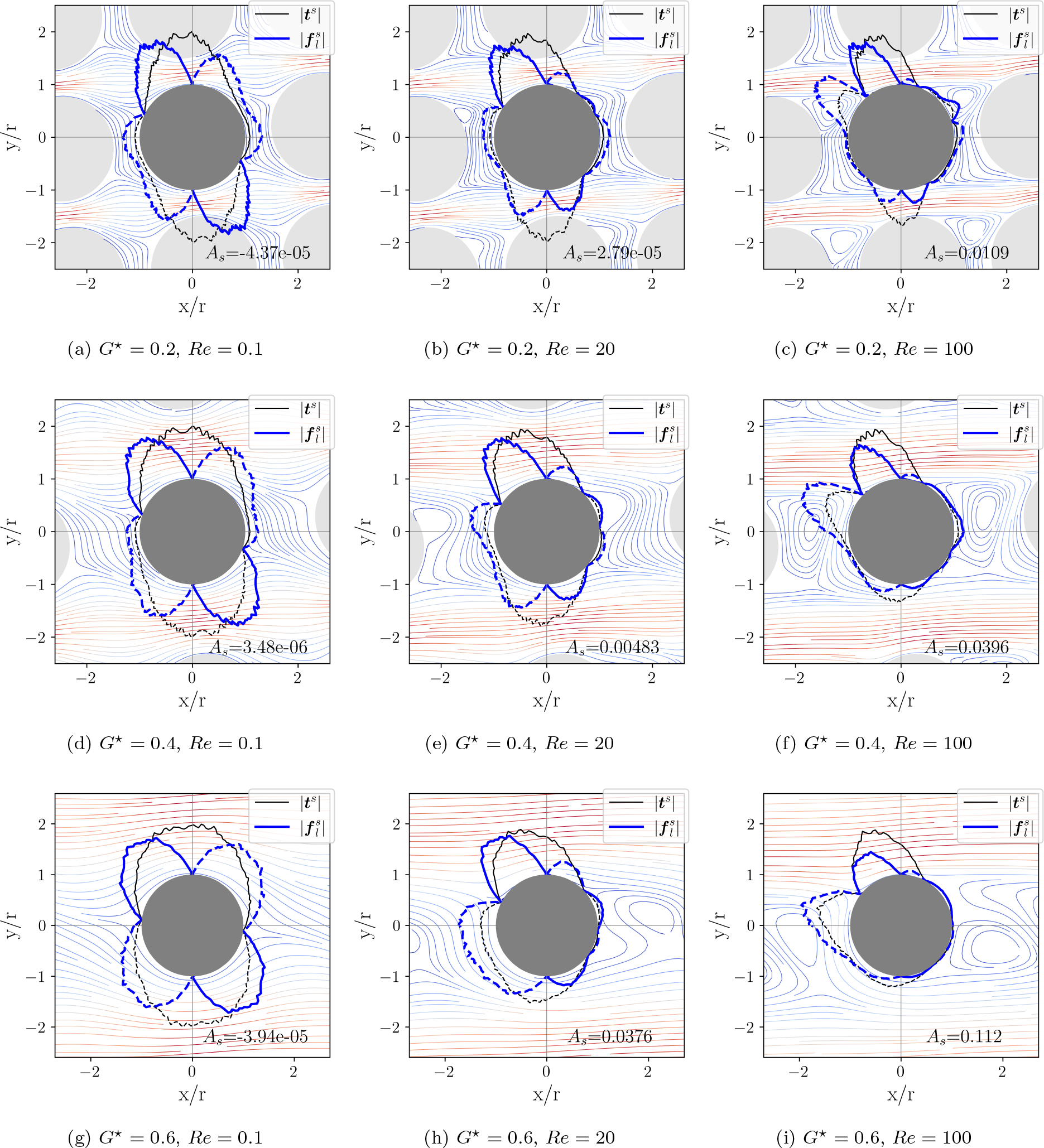
Polar representation of the shear contribution to the traction (blue) and lift (black) along the pillar surface for (a–c) *G*^⋆^ = 0.2, (d–f) *G*^⋆^ = 0.4 and (g–i) *G*^⋆^ = 0.6 at *α* = 5*°* for different values of *Re*. The average flow direction is along the *x*-axis. The traction and lift distributions are normalised by their maximum values. Positive values are shown as a solid line, negative values as a dashed line. The value of the shear contribution to the anisotropy is reported in each panel. The fluid flow is shown as streamlines using the same colouring as Fig. 3.

It can be seen from both figures that, at low *Re*, the traction and lift distributions are highly symmetric, which signifies the flow reversibility in the Stokes limit. Due to this symmetry, the surface integral over the lift distribution nearly vanishes and, thus, *A ≈* 0. Upon an increase of *Re*, the symmetry is broken in a non-trivial way, leading to a non-zero anisotropy. By definition, there is no lift generated by normal forces (Fig. 7) at the upstream and downstream points of the pillar where the surface normal is parallel to the flow direction (*x*-axis). Likewise, the lift generated by the shear forces (Fig. 8) identically vanishes at the top and bottom points of the pillar where the surface normal is parallel to the lift direction (*y*-axis).

Inspecting Fig. 7, we observe that, as *Re* increases, the normal component of the traction vector on the upstream side of the pillar becomes skewed toward the top side (from the BL to the TL quadrant). Two mechanisms contribute to this effect. First, as fluid decelerates significantly at the upstream stagnation point, pressure increases, leading to increased normal component of the traction vector on the pillar in the vicinity of the stagnation point (TL quadrant). Second, the pressure in the fluid near the BL part of the pillar decreases relatively to the pressure magnitude in the other three quadrants. Thus, the normal component of the traction vector in the BL quadrant decreases. Both mechanisms contribute negatively to lift, leading to an increase in anisotropy, according to Eq. (8).

Further inspection of Fig. 7 reveals that, with increasing *Re*, the normal component of the traction vector in the BR quadrant becomes less negative than in the TR quadrant. This change in traction distribution is caused by the skewing of the velocity profile in the vertical gap (Fig. 4), which results in a higher velocity (and therefore lower pressure) near the top side of the pillar than near the bottom side. As a consequence, the net lift force increases, contributing to a decrease in anisotropy. The two identified effects — the role of the TL stagnation point and the skewed velocity profile in the vertical gap — oppose each other but do not cancel exactly. For *Re >* 5, outside the Stokes regime, there is a net change in anisotropy which can be positive or negative, and is particularly significant for larger *G*^⋆^.

The picture is different for the contribution of the shear component of the traction vector to the lift (Fig. 8). For small *Re*, the shear component of the traction is largest around the top and bottom sides of the pillar but negligible at the upstream and downstream sides of the pillar. This observation can be explained by the main flow through the vertical gap (in positive *x*-direction) being fast (therefore creating a higher shear stress at the top and bottom sides of the pillar), while the lateral flow through the horizontal gap (in negative *y*-direction) is relatively slow (therefore creating little shear stress).

Upon increasing *Re*, the lift associated with the shear stress is primarily determined by the fluid flowing around the pillar near the stagnation point in the TL quadrant, as can be seen clearly in the right column of Fig. 8. It is helpful to consider two primary contributions to the lift in the vicinity of the stagnation point: the shear stress caused by the clockwise flow along the top side of the pillar and the shear stress caused by the counter-clockwise flow around the upstream side of the pillar. The former tends to lift the pillar upward (contributing to negative anisotropy), while the latter tends to push the pillar downward (negative lift, contributing to positive anisotropy). Importantly, the shear stress of the clockwise flow is largest in a region where the tangential vector to the pillar surface is nearly parallel to the main flow direction (*x*-axis), hence shear forces generate hardly any lift. However, the shear stress of the counter-clockwise flow is concentrated in a region where the surface tangent vector is essentially parallel to the lateral direction (*y*-axis), hence shear forces in this region contribute strongly to the lift.

For larger *G*^⋆^, we can see that the traction caused by counter-clockwise flow increases since the flow in this region is faster, and the high-speed flow region is pushed closer to the pillar surface than for smaller *G*^⋆^ (Fig. 3). Finally, since the flow in the vortex at the downstream side of the pillar is slow, there is hardly any lift generated by shear stresses in the vortex region.

In summary, we have identified several mechanisms that either increase or decrease lift and, therefore, decrease or increase anisotropy, all as function of *Re* and *G*^⋆^ (and to a lesser degree of *α*):

1. Normal component of traction vector due to upstream stagnation point (TL quadrant): 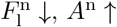
2. Normal component of traction vector due to to low pressure magnitude (BL quadrant): 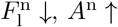
3. Normal component of traction vector due to tilted horizontal velocity profile (TR and BR quadrants): 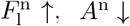
4. Shear component of traction vector due to clockwise flow along top side: 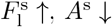
5. shear component of traction vector due to counter-clockwise flow along upstream/bottom sides: 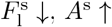

Only mechanisms 3 and 4 contribute to a decrease in anisotropy. However, mechanism 4 has been shown never to be stronger than mechanism 5, as the shear contribution to anisotropy is always positive (Fig. 6d–f)). Therefore, the net effect of mechanisms 4 and 5 always leads to an increase in anisotropy. Thus, we can conclude that the cause of the negative anisotropy for intermediate *Re* is mechanism 3, the positive lift due to pressure forces caused by the skewed velocity profile.

To further quantify the influence of the tilted horizontal velocity profile (mechanism 3) on the sign of the anisotropy, we split the contribution of the normal component of the traction vector to the lift force distribution along the pillar into regions bounded by sign changes. From Fig. 7 we see that there are generally four distinct regions: two with a negative contribution to the total lift, primarily existing in the TL and BR quadrants, and two with a positive contribution in the TR and BL quadrants. We assume that the region occupying the TL quadrant is associated with mechanism 1, and that the region occupying the BL quadrant is associated with mechanism 2. We further assume that the regions occupying the TR and BR quadrants are associated with mechanism 3. Annotations of the regions of the lift distribution with the mechanism associated with them can be seen in Fig. 7a for reference.

We integrate the lift distribution in associated with mechanisms 1 and 2 to estimate their combined effect, and we do the same for the two regions associated with mechanism 3. The results, non-dimensionalised by 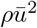, are shown in Fig. 9. Values for *Re* = 100 are not shown for *G*^⋆^ = 0.6, as the regions occupying the BL and BR quadrants merge together.

**FIG. 9.**
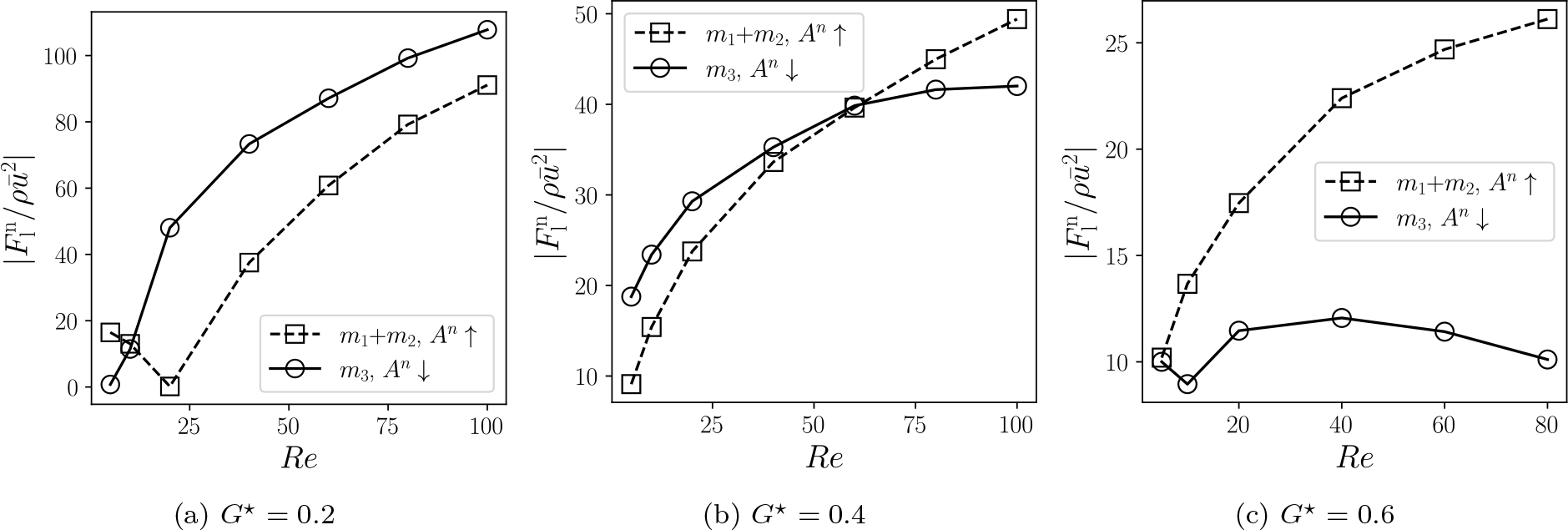
Magnitude of normal lift force associated with the different mechanisms identified in section III D. The sum of mechanisms 1 and 2 (*m*_1_ and *m*_2_), associated with the upstream stagnation point and vertical flow and contributing to positive anisotropy, is compared with mechanism 3 (*m*_3_), associated with the skewed horizontal velocity profile and contributing to negative anisotropy. When the solid line is above the broken line, we expect to see an overall negative value of the normal anisotropy, *A*n. For *G*^⋆^ = 0.6, values are only plotted up to *Re* = 80, as the lift profile has only three zeros at *Re* = 100.

Fig. 9 shows that, for *G*^⋆^ = 0.2, the mechanism associated with negative anisotropy dominates over the entire range of *Re* while both curves are close to each other, thus leading to a small negative anisotropy as found in Fig. 6(c). For *G*^⋆^ = 0.4, the mechanism associated with negative anisotropy dominates at low *Re* but begins to level off as *Re* increases, whereas the mechanisms associated with positive anisotropy increase more steadily, leading to the dominating mechanism swapping around *Re* = 60, which leads to the contribution of normal traction to the anisotropy becoming positive around *Re* = 60 as seen in Fig. 6(c). Finally, for *G*^⋆^ = 0.6, the mechanism associated with positive anisotropy dominates for all values of *Re* and increases with *Re*, which can be seen as a steep increase of the contribution of normal traction to the anisotropy in Fig. 6(c).

Although the precise mechanisms underlying the observed dependencies on *Re* and *G*^⋆^ are currently unclear, it is evident that the relative effects of mechanisms 1–3 differ based on these parameters, resulting in the observed shifts in anisotropy under varying conditions.

### E. Conclusions

Deterministic lateral displacement (DLD) is a relatively recent method for microfluidic particle separation. Although well understood in the Stokes regime, the performance of DLD devices operated with higher throughput, where inertia is not small, has not been sufficiently well characterised and analysed. Earlier work has shown that, under some conditions, the flow in DLD devices can become anisotropic, *i*.*e*., the direction of flow can deviate from that of the applied driving force. Anisotropic flow in DLD devices leads to less predictable particle behaviour and, thus, problems with particle separation.

We have investigated, *via* lattice-Boltzmann simulations, the onset and behaviour of flow anisotropy of two-dimensional rotated-square DLD arrays in the inertial regime. We show that this geometry, which exhibits no anisotropy in the Stokes regime, can have either positive or negative anisotropy, depending on the Reynolds number (*Re*), the reduced gap size (*G*^⋆^), and the array inclination angle (*α*). We find that anisotropy becomes significant for *Re >* 5, where inertial effects start to play an important role. The anisotropy increases strongly with *Re* for the largest gap size, *G*^⋆^ = 0.6, indicating that the fluid prefers to flow along the principal axis of the array. For smaller gaps, *G*^⋆^ = 0.4 and *G*^⋆^ = 0.2, anisotropy first becomes negative and only increases again at larger *Re*. We find that increasing *α* increases the magnitude of the anisotropy, but does not significantly affect the overall behaviour.

By analysing the contributions of the normal and shear components of the traction vector to the local forces acting on the pillar surface, we identify several mechanisms that either increase or decrease anisotropy, as function of *Re* and *G*^⋆^. We show that the negative contributions to the anisotropy can be attributed purely to traction forces normal to the pillar surface. We suggest that this negative contribution to the anisotropy arises from an asymmetric tilting of the velocity profile between the pillars, which leads to a pressure differential along the axis orthogonal to the flow axis.

Our findings provide valuable insights into the anisotropic behaviour of fluid flow over periodic square arrays of cylinders. Our work could be useful for the design of microfluidic devices and other applications where the control of fluid flow over periodic arrays of obstacles is important. Future work could extend this finite-inertia study to include the behaviour of particles traversing the array, to investigate three-dimensional arrays to include the effects of top and bottom walls and provide more quantitative results for real-world devices, or to investigate how anisotropy can be controlled with different post shapes at finite inertia.

## ACKNOWLEDGEMENTS

This work was supported in part by the Carnegie Trust Vacation Scholarship scheme. T.K. received funding from the European Research Council (ERC) under the European Union’s Horizon 2020 research and innovation program (No. 803553).

For the purpose of open access, the author has applied a Creative Commons Attribution (CC BY) licence to any Author Accepted Manuscript version arising from this submission.

## AUTHOR CONTRIBUTIONS

**Calum Mallorie:** Conceptualisation (equal); Formal analysis (lead); Methodology (equal); Visualisation (lead); Writing – original draft (lead); Writing – review and editing (equal). **Rohan Vernekar:** Conceptualisation (equal); Writing – review and editing (equal); **Benjamin Owen:** Formal analysis (supporting); Methodology (supporting); Writing – review and editing (equal). **Timm Krüger:** Conceptualisation (equal); Formal analysis (equal); Methodology (equal); Visualisation (supporting); Writing – original draft (supporting); Writing – review and editing (equal). **David Inglis:** Conceptualisation (equal); Writing – original draft (supporting); Writing – review and editing (equal).

## Appendix: Supplementary material

### 1. Interpolation-extrapolation scheme

**FIG. S1.**
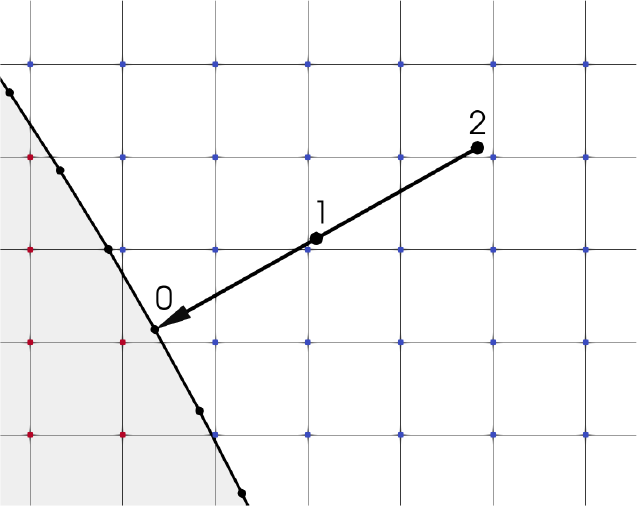
Illustration of the employed interpolation-extrapolation scheme. Lattice points are shown in red (solid nodes) and blue (fluid nodes). Flow information at points 1 and 2 is bilinearly interpolated from the four nodes defining the surrounding voxels, respectively. The interpolated values are then linearly extrapolated to point 0 on the cylinder surface.

## References

[1] Ping Hu, Wenhua Zhang, Hongbo Xin, and Glenn Deng. Single cell isolation and analysis, volume 4. Frontiers Media S.A., 2016.

[2] L. R. Huang, E. C. Cox, R. H. Austin, and J. C. Sturm. Continuous particle separation through deterministic lateral displacement. Science, 304(5673):987–990, May 2004.

[3] Curt I. Civin, Tony Ward, Alison M. Skelley, Khushroo Gandhi, Zendra Peilun Lee, Christopher R. Dosier, Joseph L. D’Silva, Yu Chen, MinJung Kim, James Moynihan, Xiaochun Chen, Lee Aurich, Sergei Gulnik, George C. Brittain, Diether J. Recktenwald, Robert H. Austin, and James C. Sturm. Automated leukocyte processing by microfluidic deterministic lateral displacement. Cytometry Part A, 89(12):1073–1083, 2016. eprint: https://onlinelibrary.wiley.com/doi/pdf/10.1002/cyto.a.23019.

[4] Kerwin Kwek Zeming, Rohan Vernekar, Mui Teng Chua, Kai Yun Quek, Greg Sutton, Timm Krüger, Win Sen Kuan, and Jongyoon Han. Label-Free Biophysical Markers from Whole Blood Microfluidic Immune Profiling Reveal Severe Immune Response Signatures. Small, 17(12):2006123, March 2021. Publisher: John Wiley and Sons Inc.

[5] D. W. Inglis, J. A. Davis, R. H. Austin, and J. C. Sturm. Critical particle size for fractionation by deterministic lateral displacement. Lab on a Chip, 6(5):655–658, 2006.

[6] David W Inglis, Nick Herman, and Graham Vesey. Highly accurate deterministic lateral displacement device and its application to purification of fungal spores. Biomicrofluidics, 4(2), 2010.

[7] Haakan N. Joensson, Mathias Uhlén, and Helene Andersson Svahn. Droplet size based separation by deterministic lateral displacement- separating droplets by cell-induced shrinking. Lab on a Chip, 11(7):1305–1310, 2011.

[8] John Alan Davis. Microfluidic Separation of Blood Components through Deterministic Lateral Displacement. Thesis, Princeton University, Princeton New Jersey, 2008.

[9] J. McGrath, M. Jimenez, and H. Bridle. Deterministic lateral displacement for particle separation: a review. Lab on a Chip, 14(21):4139–4158, 2014.

[10] Maike S. Wullenweber, Jonathan Kottmeier, Ingo Kampen, Andreas Dietzel, and Arno Kwade. Numerical Study on High Throughput and High Solid Particle Separation in Deterministic Lateral Displacement Microarrays. Processes, 11(8):2438, August 2023. Number: 8 Publisher: Multidisciplinary Digital Publishing Institute.

[11] Arian Aghilinejad, Mohammad Aghaamoo, and Xiaolin Chen. On the transport of particles/cells in high-throughput deterministic lateral displacement devices: Implications for circulating tumor cell separation. Biomicrofluidics, 13:34112, 2019.

[12] Brian M. Dincau, Arian Aghilinejad, Taylor Hammersley, Xiaolin Chen, and Jong-Hoon Kim. Deterministic lateral displacement (DLD) in the high Reynolds number regime: high-throughput and dynamic separation characteristics. Microfluidics and Nanofluidics, 22(6):59, 2018.

[13] Y. S. Lubbersen, M. A. I. Schutyser, and R. M. Boom. Suspension separation with deterministic ratchets at moderate Reynolds numbers. Chemical Engineering Science, 73:314–320, May 2012.

[14] Timothy J. Bowman, German Drazer, and Joelle Frechette. Inertia and scaling in deterministic lateral displacement. Biomicrofluidics, 7(6):064111, 2013.

[15] S. R. Reinecke, S. Blahout, T. Rosemann, B. Kravets, M. Wullenweber, A. Kwade, J. Hussong, and H. Kruggel-Emden. DEM-LBM simulation of multidimensional fractionation by size and density through deterministic lateral displacement at various Reynolds numbers. Powder Technology, 385:418–433, June 2021.

[16] Brian M. Dincau, Arian Aghilinejad, Xiaolin Chen, Se Youn Moon, and Jong-Hoon Kim. Vortex-free high-Reynolds deterministic lateral displacement (DLD) via airfoil pillars. Microfluidics and Nanofluidics, 22(12):137, 2018.

[17] Y.S. Lubbersen, J.P. Dijkshoorn, M.A.I. Schutyser, and R.M. Boom. Visualization of inertial flow in deterministic ratchets. Separation and Purification Technology, 109:33–39, 2013.

[18] T. Kulrattanarak, R. G. M. van der Sman, Y. S. Lubbersen, C. G. P. H. Schroën, H. T. M. Pham, P. M. Sarro, and R. M. Boom. Mixed motion in deterministic ratchets due to anisotropic permeability. Journal of Colloid and Interface Science, 354(1):7–14, February 2011.

[19] Sung-Cheol Kim, Benjamin H. Wunsch, Huan Hu, Joshua T. Smith, Robert H. Austin, and Gustavo Stolovitzky. Broken flow symmetry explains the dynamics of small particles in deterministic lateral displacement arrays. Proceedings of the National Academy of Sciences, 114(26):E5034–E5041, June 2017. Publisher: Proceedings of the National Academy of Sciences.

[20] Rohan Vernekar, Timm Krüger, Kevin Loutherback, Keith Morton, and David W. Inglis. Anisotropic permeability in deterministic lateral displacement arrays. Lab on a Chip, 17(19):3318–3330, September 2017.

[21] Axel Hochstetter, Rohan Vernekar, Robert H. Austin, Holger Becker, Jason P. Beech, Dmitry A. Fedosov, Gerhard Gompper, Sung-Cheol Kim, Joshua T. Smith, Gustavo Stolovitzky, Jonas O. Tegenfeldt, Benjamin H. Wunsch, Kerwin K. Zeming, Timm Krüger, and David W. Inglis. Deterministic Lateral Displacement: Challenges and Perspectives. ACS Nano, 14(9):10784–10795, September 2020. Publisher: American Chemical Society.

[22] Donald L. Koch and Anthony J. C. Ladd. Moderate Reynolds number flows through periodic and random arrays of aligned cylinders. Journal of Fluid Mechanics, 349:31–66, October 1997.

[23] A. Dybbs and R. V. Edwards. A New Look at Porous Media Fluid Mechanics — Darcy to Turbulent. In Jacob Bear and M. Yavuz Corapcioglu, editors, Fundamentals of Transport Phenomena in Porous Media, NATO ASI Series, pages 199–256. Springer Netherlands, Dordrecht, 1984.

[24] Michel Quintard and Stephen Whitaker. Transport in ordered and disordered porous media: volume-averaged equations, closure problems, and comparison with experiment. Chemical Engineering Science, 48(14):2537–2564, July 1993.

[25] Dilip Natarajan and Trung Van Nguyen. A Two-Dimensional, Two-Phase, Multicomponent, Transient Model for the Cathode of a Proton Exchange Membrane Fuel Cell Using Conventional Gas Distributors. Journal of The Electrochemical Society, 148(12):A1324, November 2001. Publisher: IOP Publishing.

[26] Hanieh Bazyar, Pengyu Lv, Jeffery A. Wood, Slawomir Porada, Detlef Lohse, and Rob G. H. Lammertink. Liquid–liquid displacement in slippery liquid-infused membranes (SLIMs). Soft Matter, 14(10):1780–1788, March 2018. Publisher: The Royal Society of Chemistry.

[27] Manish Kumar, Jeffrey S. Guasto, and Arezoo M. Ardekani. Transport of complex and active fluids in porous mediaa). Journal of Rheology, 66(2):375–397, March 2022.

[28] Allen G. Hunt and Muhammad Sahimi. Flow, Transport, and Reaction in Porous Media: Percolation Scaling, Critical-Path Analysis, and Effective Medium Approximation. Reviews of Geophysics, 55(4):993–1078, 2017. eprint: https://onlinelibrary.wiley.com/doi/pdf/10.1002/2017RG000558.

[29] A. Tamayol, A. Khosla, B. L. Gray, and M. Bahrami. Creeping flow through ordered arrays of micro-cylinders embedded in a rectangular minichannel. International Journal of Heat and Mass Transfer, 55(15):3900–3908, July 2012.

[30] Tobias O. M. Forslund, I. A. Sofia Larsson, Henrik Lycksam, J. Gunnar I. Hellström, and T. Staffan Lundstrom. Non-Stokesian flow through ordered thin porous media imaged by tomographic-PIV. Experiments in Fluids, 62(3):46, March 2021.

[31] T. O. M. Forslund, I. A. S. Larsson, J. G. I. Hellström, and T. S. Lundström. Steady-State Transitions in Ordered Porous Media. Transport in Porous Media, 149(2):551–577, September 2023.

[32] A. Xu, T. S. Zhao, L. Shi, and J. B. Xu. Lattice Boltzmann Simulation of Mass Transfer Coefficients for Chemically Reactive Flows in Porous Media. Journal of Heat Transfer, 140(052601), January 2018.

[33] D. Lasseux, A. A. Abbasian Arani, and A. Ahmadi. On the stationary macroscopic inertial effects for one phase flow in ordered and disordered porous media. Physics of Fluids, 23(7):073103, July 2011.

[34] Thejas Hulikal Chakrapani, Hanieh Bazyar, Rob G. H. Lammertink, Stefan Luding, and Wouter K. den Otter. The permeability of pillar arrays in microfluidic devices: an application of Brinkman’s theory towards wall friction. Soft Matter, December 2022. Publisher: The Royal Society of Chemistry.

[35] Kawkab Ahasan, Christopher M. Landry, Xiaolin Chen, and Jong-Hoon Kim. Effect of angle-of-attacks on deterministic lateral displacement (DLD) with symmetric airfoil pillars. Biomedical Microdevices, 22(2):42, June 2020.

[36] Timm Krüger, Halim Kusumaatmaja, Alexandr Kuzmin, Orest Shardt, Goncalo Silva, and Erlend Magnus Viggen. The Lattice Boltzmann Method: Principles and Practice. Graduate Texts in Physics. Springer International Publishing, 2017.

[37] S. Succi. The lattice Boltzmann equation for fluid dynamics and beyond. Clarendon Press, 2001.

[38] P. L. Bhatnagar, E. P. Gross, and M. Krook. A Model for Collision Processes in Gases. I. Small Amplitude Processes in Charged and Neutral One-Component Systems. Physical Review, 94(3):511–525, May 1954. Publisher: American Physical Society.

[39] Zhaoli Guo, Chuguang Zheng, and Baochang Shi. Discrete lattice effects on the forcing term in the lattice Boltzmann method. Physical Review E, 65(4):046308, April 2002. Publisher: American Physical Society.

[40] Anthony J. C. Ladd. Numerical simulations of particulate suspensions via a discretized Boltzmann equation. Part 1. Theoretical foundation. Journal of Fluid Mechanics, 271:285–309, July 1994. Publisher: Cambridge University Press.

